# Both functional trait divergence and trait plasticity confer polyploid advantage in changing environments

**DOI:** 10.1101/274399

**Authors:** Na Wei, Richard Cronn, Aaron Liston, Tia-Lynn Ashman

## Abstract

Polyploidy, or whole genome duplication, exists in all eukaryotes and is thought to drive ecological and evolutionary success especially in plants. The mechanisms of polyploid success in ecologically relevant contexts, however, remain largely unknown. Here we conducted an extensive test of functional trait divergence and trait plasticity in conferring polyploid fitness advantage in changing environments by growing clonal replicates of a worldwide genotype collection of six polyploid and five diploid wild strawberry (*Fragaria*) species in three climatically different common gardens. Among leaf functional traits, we detected divergence in means but not plasticities between polyploids and diploids, suggesting that increased genomic redundancy does not necessarily translate into broader phenotypic amplitude in polyploids. Across the heterogeneous garden environments, however, polyploids exhibited fitness advantage, which was conferred by both trait means and adaptive trait plasticities, supporting a ‘jack-and-master’ hypothesis for polyploids. Our findings provide unparalleled insight into the prevalence and persistence of polyploidization.

## INTRODUCTION

Polyploidy (or whole-genome duplication) enlarges and diversifies an organism’s genome, with profound influence on phenotype and fitness (Otto & Whitton 2000; Ramsey & Ramsey 2014; Soltis *et al.* 2016). While polyploidy exists in all eukaryotes, some of the best-known examples of polyploids in angiosperms are among crops (Renny-Byfield & Wendel 2014) and invasive plants (te Beest *et al.* 2012), and the repeated and pervasive occurrence of polyploidy throughout the plant kingdom reflects its widespread adaptive significance (Van de Peer *et al.* 2017). Despite its evolutionary importance, the mechanisms of polyploid advantage are largely unknown. A leading, yet rarely tested, hypothesis is that polyploid fitness advantage arises from enhanced means in functional traits and/or the ability to adjust phenotype (i.e. trait plasticity) in response to environmental change (Levin 1983; Van de Peer *et al.* 2017).

Polyploidy can alter plant phenotype (Levin 1983; Soltis *et al.* 2014). Phenotypic variation at the cellular level (e.g. an increase in cell size) as a result of an increase in ploidy was first recognized in early cytological studies of synthetic polyploids (reviewed in Ramsey & Ramsey 2014). This positive correlation between genome size and cell size holds across angiosperm lineages (Masterson 1994; Beaulieu *et al.* 2008), whereas for phenotype at higher levels (e.g. tissue, organism), the nucleotypic effects of genome size are shown to be weaker or absent (Knight & Beaulieu 2008). In addition to the genome size effect, polyploidy also diversifies a plant genome by merging multiple copies of genes from the same (autopolyploidy) or different species (allopolyploidy), which can have important implications for phenotype (Chen 2010; Soltis *et al.* 2014). Comparisons of phenotypic divergence between diploids and naturally occurring polyploids, in ecologically relevant contexts, have been primarily conducted in autopolyploids with intraspecific ploidal variation (Ramsey & Ramsey 2014), and have yielded mixed, and often system-specific, conclusions (e.g. Li *et al.* 1996; Maherali *et al.* 2009; Balao *et al.* 2011; Li *et al.* 2012; Hao *et al.* 2013). Moreover, the phenotypic consequences of allopolyploidy – which generates diverse genetic backgrounds and the potential to express transgressive phenotypes relative to autopolyploidy (Chen 2010) – remains unclear for most polyploid taxa (but see Manzaneda *et al.* 2015; Leal-Bertioli *et al.* 2017).

Polyploidy has the potential to alter phenotypic plasticity (Stebbins 1971), owing to genomic redundancy and plasticity (Adams & Wendel 2005; Leitch & Leitch 2008; Jackson & Chen 2010; Madlung & Wendel 2013). Relative to diploids, polyploids can potentially employ alternative copies of duplicated genes gained from diverse and possibly adaptive genetic backgrounds to respond to novel environments (Bardil *et al.* 2011; Dong & Adams 2011; Shimizu-Inatsugi *et al.* 2017). Thus, it is hypothesized that polyploids can exhibit higher phenotypic plasticity than diploids in response to changing environments. Previous work has primarily emphasized the genomic plasticity of polyploids (reviewed in Soltis *et al.* 2016), and as a result the questions of whether genome duplication translates into increased phenotype plasticity in the wild (Madlung 2013), and how phenotypic plasticity differs between diploids and polyploids (Hahn *et al.* 2012; Sánchez Vilas & Pannell 2017) remain.

Polyploidy has been demonstrated to provide selective advantages to plants under environmental stresses and instabilities (Chao *et al.* 2013; Yang *et al.* 2014; Van de Peer *et al.* 2017). However, it remains controversial whether such polyploid fitness advantage occurs only in a particular environment or can be maintained consistently across environments (Ramsey 2011; Madlung 2013; McIntyre & Strauss 2017). Several competing adaptive hypotheses have been proposed. First, elevated genetic heterozygosity and genomic plasticity may enable polyploids to occupy broader ecological niches (i.e. higher ecological amplitude) than diploids. As a result of possessing such ‘general purpose’ genotypes (Stebbins 1971), polyploids could exhibit high fitness and fitness homoeostasis in heterogeneous environments (i.e. high intercept and low slope in a fitness reaction norm; ‘jack-of-all-trades’) (Richards *et al.* 2006).

Alternatively, in the absence of fitness homoeostasis, polyploids may still maintain higher fitness than diploids across a broad range of environments. This fitness strategy (manifesting as high intercept and high slope) can be referred to as ‘jack-and-master’, following the usage previously proposed for plant invasion (Richards *et al.* 2006), but not considering slope difference between polyploids and diploids. Lastly, polyploids and diploids may both be habitat specialists, exhibiting high fitness in alternative environments (i.e. ‘master-of-some’). While these adaptive hypotheses have been broadly tested among invasive and native plant species (e.g. Richards *et al.* 2006; Davidson *et al.* 2011), their evaluations with respect to polyploidy are not only limited to a few intra-and interspecific systems (Petit & Thompson 1997; Bretagnolle & Thompson 2001; McIntyre & Strauss 2017), but more importantly they lack the mechanisms that connect the fitness of diploids and polyploids to functional traits and trait plasticity.

In this study, we take advantage of the fact that polyploidy is an important mode of speciation in wild strawberries (*Fragaria*), a genus with a broad distribution in the Northern Hemisphere (Staudt 1999; Liston *et al.* 2014). By growing clonal replicates of a worldwide collection of *Fragaria* genotypes of six allopolyploid and five diploid species in three climatically different common gardens in Oregon, USA, we addressed the following questions: (1) Do functional traits differ between diploids and polyploids? (2) Do polyploids demonstrate higher trait plasticity than diploids in response to environmental change? (3) Is there a polyploid fitness advantage across diverse garden environments? (4) If so, then is the polyploid fitness advantage conferred by trait means, trait plasticities, or both? Our results indicate that polyploids and diploids differ in several leaf functional traits. Although different functional traits (Table 1) display varying degrees of plasticity, trait plasticity is of similar magnitude between diploids and polyploids. Furthermore, our study demonstrates that polyploids have a fitness advantage across all experimental environments, which is gained through both differences in trait means and adaptive plasticities.

**Table 1:** Key variables of the common garden experiment.

|  | Description | Units | Function<sup>†</sup> |
| --- | --- | --- | --- |
| **Leaf functional traits** |  |  |  |
| Specific leaf area (SLA) | Light-capturing leaf area per unit dry mass | mm<sup>2</sup> mg<sup>-1</sup> | SLA reflects the thickness and/or dry mass content of leaf tissue. High SLA permits high leaf carbon gain. |
| Stomatal density (SD) | Number of abaxial stomata per unit leaf area | mm<sup>-2</sup> | SD regulates CO<sub>2</sub> intake and water transpiration, reflecting the trade-off between gas conductance and epidermal construction cost. |
| Stomatal length (SL) | Abaxial guard cell length | μm | SL correlates negatively with SD. |
| Vein density (VLA) | Total minor vein length per unit leaf area | mm mm<sup>-2</sup> | VLA supports leaf hydraulic conductance. Low VLA however reduces construction cost. |
| Trichome density (TD) | Total ab- and adaxial trichomes per unit leaf area | mm<sup>-2</sup> | TD influences the ability of plants to prevent water loss. |
| Nitrogen content ($N_{\text{mass}}$) | Leaf nitrogen per unit dry mass | % | $N_{\text{mass}}$, required for photosynthetic proteins, supports leaf photosynthetic potential. |
| Carbon isotope discrimination ($\Delta^{13}\text{C}$) | Amount of isotope discrimination against <sup>13</sup>C relative to <sup>12</sup>C during photosynthesis | ‰ | $\Delta^{13}\text{C}$ indicates photosynthetic water use efficiency, integrated over the life span of a leaf. Low $\Delta^{13}\text{C}$ reflects high water use efficiency. |
| **Plant fitness proxies** |  |  |  |
| Survival | Presence (1) or absence (0) of a plant |  |  |
| Plant size | LN × LA | dm<sup>2</sup> |  |
| Stolon mass | Dry mass of stolons | g |  |
| **Leaf number and size** |  |  |  |
| Leaf number (LN) | Number of leaves per plant |  |  |
| Leaf area (LA) | Area of the largest leaf | dm<sup>2</sup> |  |
| Central leaflet width (CLW) | Width of the central leaflet of the largest leaf | mm |  |

†References for each trait function: SLA (Poorter *et al.* 2009); SD and SL (Hetherington & Woodward 2003); VLA (Sack & Scoffoni 2013); TD (Ehleringer & Björkman 1978; Sletvold & Agren 2012); $N_{\text{mass}}$ (Wright *et al.* 2004); $\Delta^{13}\text{C}$ (Farquhar & Richards 1984).

## MATERIALS AND METHODS

### Study system and field collection

Wild strawberries (*Fragaria*) are perennial herbaceous plants that reproduce both sexually by seed and asexually by plantlets on stolons (Staudt 1999). *Fragaria* originated around 3–8 Mya (Liston *et al.* 2014; Qiao *et al.* 2016), and has 22 extant species, half of which are polyploids (Staudt 1999; Liston *et al.* 2014). *Fragaria* have two centers of species diversification (in East Asia and Europe–North America) (Liston *et al.* 2014) and occupy diverse ecological habitats (Staudt 1999; Johnson *et al.* 2014). For this study, we considered diploid and polyploid *Fragaria* that occur in North America, South America, Europe and Japan (Fig. 1). The six allopolyploid strawberries are hexaploid (6*x*) *F*. *moschata*, octoploid (8*x*) *F*. *chiloensis* ssp. *pacifica, F*. *chiloensis* ssp. *chiloensis, F*. *virginiana* ssp. *platypetala, F*. *virginiana* ssp. *virginiana*, and decaploid (10*x*) *F*. *cascadensis*. The five diploid strawberries are *F*. *vesca* ssp. *bracteata, F*. *vesca* ssp. *americana, F*. *vesca* ssp. *vesca, F*. *viridis*, and *F*. *iinumae*. Our previous studies of polyploid *Fragaria* genomes (Tennessen *et al.* 2014; Kamneva *et al.* 2017; Wei *et al.* 2017; Dillenberger *et al.* 2018) have indicated the repeated and independent events of allopolyploid speciation for the aforementioned polyploids: the 10*x* and 8*x* taxa are derived from the 2*x F*. *vesca* ssp. *bracteata* and *F*. *iinumae*, and the 6*x* species is derived from the 2*x F*. *viridis* and *F*. *vesca* ssp. *vesca*. Details of the worldwide collection of *Fragaria* are available on our Wild Strawberry website (http://wildstrawberry.org/; see also Data Availability).

**Figure 1:**
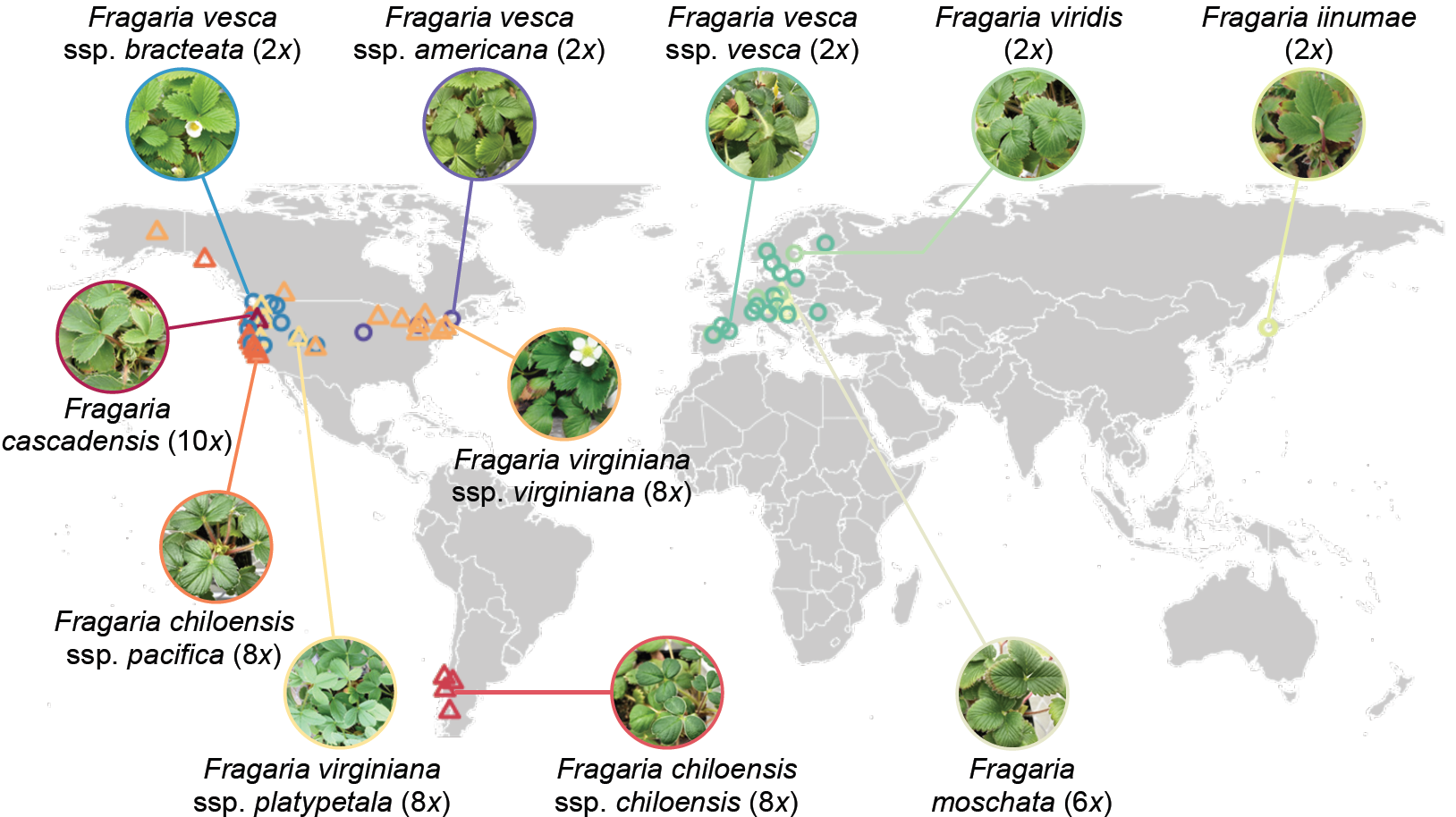
Source populations of *Fragaria* diploids and polyploids. The ploidal variation of polyploids range from hexaploid (6*x*) to octoploid (8*x*) and decaploid (10*x*). Polyploids are denoted as triangles and diploids as circles.

### Genotype and clone cultivation

In April 2015, we germinated and grew four genotypes (i.e. each from a single, open-pollinated seed of a distinct wild plant) from each of 72 total populations across the 11 species (Fig. 1; 10 populations of <4 genotypes, Table S1), in a glasshouse at the University of Pittsburgh following standard protocols (Wei *et al.* 2017). In September 2015, we harvested 12 plantlets (clones) from stolons of each of the 269 genotypes (24 genotypes had <12 clones, Table S1). Plantlets (*N* = 3137) were sent to Oregon State University (Corvallis, OR, USA), kept in dark at 16°C for one week to stimulate root growth, and then transplanted to 107 cm^2^ cone-tainers (Stuewe & Sons Inc., Tangent, OR, USA) filled with Sunshine Mix #4 soil (Sun Gro Horticulture, Agawam, MA, USA). Plantlets were grown at 18°C under natural lighting in a glasshouse for three weeks, and moved outside for one week prior to transplanting in common gardens during the Fall (October 28–November 15, 2015). At transplanting, clones had 1–2 leaves, most of which senesced over winter.

### Common gardens

Three common gardens were located in Oregon, USA: cool/coastal ‘Newport’ (44.62046°N, 124.04410°W; altitude, 5 m), temperate/valley ‘Corvallis’ (44.56107°N, 123.28911°W; 70 m) and arid/montane ‘Bend’ (44.08895°N, 121.26192°W; 1063 m), each differing in temperature and precipitation (Fig. S1). At each location, we established four raised wooden beds (18 m × 1.5 m; Fig. S1), filled with soil derived from local sources (Appendix S1; Table S2).

Plants were arranged in a complete randomized block design with ˜25 cm spacing, and one clone per genotype was randomly assigned a position in one of the four beds (blocks) at each garden location (Fig. S1). For the 24 genotypes with <12 clones, we distributed available clones evenly across garden locations, but within each garden we prioritized filling beds 1 and 2 to have at least two complete blocks each location. Empty positions (*N* = 319) were filled with non-experimental clones, which were cultivated in the same manner as the others, to maintain even plant spacing and density. Throughout the course of the experiment (October 2015–July 2016), plants received only natural precipitation at Newport and Corvallis, which reached a total of 138.5 cm and 95.5 cm, respectively (Fig. S1); however, at Bend (natural precipitation of 58.2 cm), plants were given supplemental water totaling 14.2 cm during the months of near-zero rainfall (February–April 2016; Fig. S1). All beds were protected from large herbivores using polypropylene mesh (1.6 cm × 1.6 cm) netting. Beds at Bend received straw cover (November 2015–February 2016) to minimize winter freeze damage to plant crowns.

### Functional traits and fitness proxies

We assessed a suite of leaf functional traits that reflect essential plant ecophysiological processes (Table 1) in May 2016 on experimental plants in beds 1 and 2 of each garden (*N* = 1429). Within selected beds, we counted the number of leaves (LN) of individual plants, and collected the largest leaf to measure leaf area (LA) and seven functional traits as described in Appendix S2. Among these traits, vein density (or vein length per unit area, VLA) and trichome density (TD) were measured only at Corvallis and Bend (*N* = 950), as leaves of plants at Newport were too small for additional measures. Leaf nitrogen content (*N*_mass_) and carbon isotope discrimination (Δ^13^C) were obtained for a subset of randomly chosen genotypes per population at individual gardens (*N* = 210).

As most plants did not flower in 2016, we estimated plant fitness based on survival, growth (i.e. plant size) and asexual reproduction (i.e. stolon mass). We scored plant survival in May 2016 on plants in all four beds in each garden. For plants in beds 1 and 2 of each garden, we estimated plant size as the product of LN and LA (Table 1). All experimental plants survived to the time (July 7–14, 2016) when we harvested stolons, which were dried at 65°C for one week prior to weighing.

### Climatic niche distance

Plant functional traits and fitness can be influenced by climatic differences between source populations and experimental gardens, or the ‘climatic niche distance’ (CND). To estimate CND, we extracted the 19 bioclimatic variables (current conditions, 1970–2000) at 30-arcsec resolution (or 2.5-arcmin resolution for west coast populations of North America), as well as altitude estimates, from WorldClim v2.0 (Fick & Hijmans 2017) for the 72 source populations and the three garden locations. We conducted a principal component analysis of these 20 variables using prcomp() in R v3.3.3 (R Core Team 2017). The first five principal components (Fig. S2), accounting for 94.2% of the variation, were used to calculate the Euclidean CND between each source population and each garden using the R package pdist (Wong 2013).

We did not include soil parameters in CND estimation because topsoil (0–30 cm) data were missing or incomplete for 14 *Fragaria* populations from the Harmonized World Soil Database (HWSD, FAO/IIASA/ISRIC/ISS-CAS/JRC 2012), and the soil variables measured for the raised beds were not available from HWSD.

### Data analyses

To evaluate whether diploids and polyploids differ in functional trait means, we performed general linear mixed models (LMMs) using the package lme4 (Bates *et al.* 2015). The response variables of LMMs considered genotypic values of each trait (i.e. the average of two clones) at each garden. We applied power transformation to response variables using the Box–Cox method in the R package car (Fox & Weisberg 2011) to improve normality. The fixed effects of LMMs included ploidy level (i.e. diploid or polyploid), garden, CND and the interactions of ploidy level with the latter two predictors, as well as central leaflet width (Table 1). We incorporated central leaflet width to account for functional trait variation potentially attributable to leaf characteristics (e.g. stage, expansion, vigor) rather than ploidy levels. The random effects included populations nested within species and then within ploidy levels. For the main effects of predictors and the interactions, the least-squares means were estimated using the R package lmerTest (Kuznetsova *et al.* 2016), and the statistical significance was evaluated by Type III sums of squares.

To evaluate whether polyploids express higher trait plasticity than diploids, we estimated plasticity for each trait and genotype using relative distance plasticity index (RDPI) and phenotypic plasticity index (PI) (Valladares *et al.* 2006). For traits that were only measured at two gardens (vein density and trichome density), plasticity was calculated as trait distance (in absolute value) of the same genotype between the two environments, divided by the mean (for RDPI) or by the maximum (for PI) of the two genotypic trait values. For traits measured at all three gardens, RDPI and PI were calculated as the mean of the three pairwise distances. Trait plasticity, power transformed if necessary, was taken as the response variable in a LMM with nested random effects for each trait. The fixed effects included ploidy level and CND, and their interactions.

To evaluate whether polyploids have higher fitness than diploids, we estimated genotypic fitness at each garden using a composite fitness index as the product of genotypic survival rate, plant size and stolon mass. The genotypic survival rate was calculated as the proportion of clones that survived to May 2016 in all four beds per garden. The genotypic plant size and stolon mass were the average of the two clones measured per garden. As many plants produced zero stolons at Newport, we adjusted the genotypic stolon mass at each garden by adding 0.01 g. Our estimate of fitness represented a relatively equal contribution from each of the three components, owing to their similar scales across gardens (i.e. survival rate, median = 1; plant size, 0.64 dm^2^; stolon mass, 0.56 g). Fitness after transformation (with the power parameter λ = 0.1) was taken as the response variable in a LMM with nested random effects, in which the fixed effects included ploidy level, garden and CND, and the interactions of ploidy level with the other two predictors.

To determine whether plant fitness over all garden environments is conferred by trait means or plasticities, we used mean fitness per genotype across the three gardens as the response variable (with power transformation, λ = 0.1) in LMMs with nested random effects for individual traits (except vein density and trichome density with mean fitness of two gardens). The fixed effects of a LMM included the main effects of trait mean and trait plasticity and their interactions with ploidy level, as well as CND mean. Here trait mean and CND mean were defined as the genotypic trait and CND, respectively, averaged over the three gardens (or two gardens for vein density and trichome density). LMMs with trait plasticities of RDPI and PI were performed separately, but as they yielded similar patterns we only reported the results based on RDPI. To compare the magnitude of the respective effects of trait mean and trait plasticity on average fitness, we reported the standardized coefficients (β’) of these fixed effects using the R package sjPlot (Lüdecke 2017).

Our analyses did not control for phylogenetic relatedness among these *Fragaria* species for two reasons. First, polyploid phylogenies are reticulate and complex especially for allopolyploids (Wei *et al.* 2017), and thus their evolutionary histories cannot be accurately represented by a bifurcating tree (e.g. chloroplast tree). Second, phylogenetically informed approaches rest on the assumption that differences between ploidy levels are influenced by separate evolutionary trajectories of diploid and polyploid lineages. In fact, as mentioned above, some polyploid and diploid *Fragaria* are more closely related to each other, relative to species within the same ploidy level. Thus, comparing multiple diploids and polyploids of independent and diverse origins as two separate groups can broadly inform the ecological consequences of polyploidy.

## Results

### Do functional traits differ between diploids and polyploids?

Diploid and polyploid *Fragaria* differed in most leaf functional traits (Fig. 2; Table S3), either consistently across environments (e.g. stomatal length and vein density) or in certain environments (e.g. SLA, stomatal density and nitrogen content).

**Figure 2:**
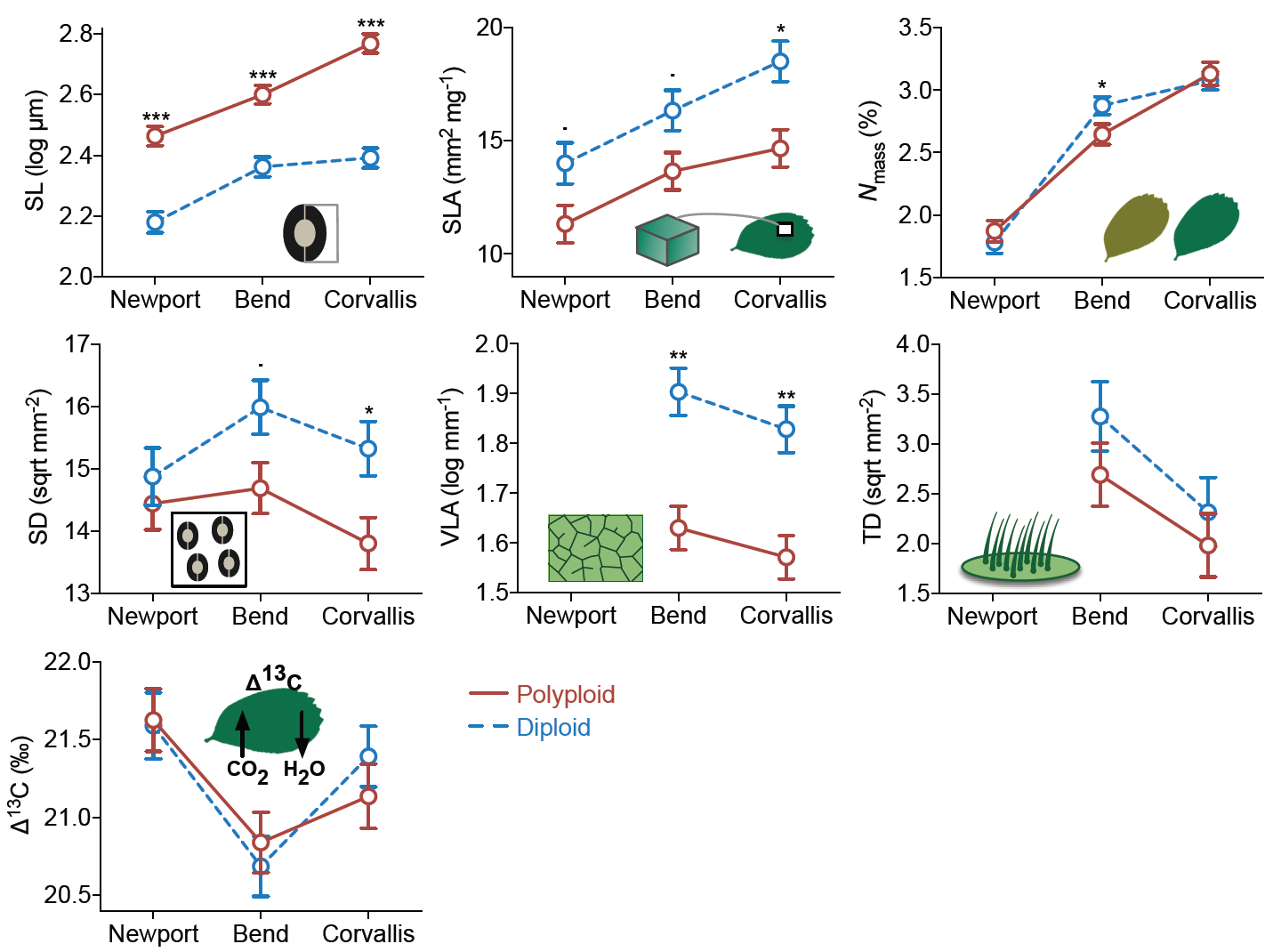
Differences in the least-squares means of leaf functional traits between diploids and polyploids. The response variables were power transformed in the general linear mixed models (see Methods). The *x*-axis is arranged from the least favorable, cool/coastal garden at Newport to the most favorable, temperate/valley garden at Corvallis. Error bars represent one standard error from the least-squares means. The dashed lines depict diploids, and solid lines depict polyploids. SL, stomatal length; SLA, specific leaf area; *N*_mass_, nitrogen content; SD, stomatal density; VLA, vein density; TD, trichome density; Δ^13^C, carbon isotope discrimination. Significance levels: ^***^, *P* < 0.001; ^**^, *P* < 0.01; ^*^, *P* < 0.05; ?, *P* = 0.053.

Polyploids possessed larger stomata than diploids in all environments (*F* = 37.56, df = 1, *P* < 0.001), while accounting for CND and central leaflet width. In the favorable environment at Corvallis, polyploids produced not only larger stomata (*t* = 8.48, *P* < 0.001) but also fewer stomata per unit leaf area (*t* = -2.57, *P* = 0.028) than diploids, which may lower the epidermal construction cost of stomata per unit area for gas exchange (de Boer *et al.* 2016) in polyploids. The general trade-off between stomatal length and density seen across vascular plants (Franks & Beerling 2009) was, nevertheless, decoupled in the stressful environment at Newport (Fig. 2); reduced stomatal length was not accompanied by increased stomatal density, for polyploids and especially diploids, which could negatively affect photosynthetic potential (Tanaka *et al.* 2013). Moreover, polyploids and diploids also differed in vein density across environments (*F* = 4.74, df = 1, *P* = 0.037; Fig. 2), with polyploids producing lower minor vein length per unit leaf area (i.e. lower hydraulic construction cost) (Sack & Scoffoni 2013).

Although the main effect of ploidy level across gardens did not influence SLA (*F* = 3.10, df = 1, *P* = 0.100; Table S3) and nitrogen content (*F* = 0.57, df = 1, *P* = 0.453), polyploids produced foliage with significantly smaller SLA than diploids at Corvallis (*t* = -3.18, *P* = 0.011), and significantly lower nitrogen content at Bend (*t* = -2.12, *P* = 0.040). In contrast, polyploids and diploids were similar in leaf traits that influence water loss (trichome density, *F* = 0.96, df = 1, *P* = 0.346) and water use efficiency (Δ^13^C, *F* = 1.46, df = 1, *P* = 0.233) in all environments (Fig. 2).

### Do polyploids demonstrate higher trait plasticity than diploids in response to environmental change?

*Fragaria* genotypes expressed plasticity for the measured traits in response to experimental environments (Fig. 2), as demonstrated by the significant main effect of garden on each trait (all *P* < 0.001; Table S3), after accounting for the influence of ploidy level, CND and central leaflet width. Quantifying plasticity using RDPI and PI yielded similar patterns in degrees of plasticity among traits: (1) Δ^13^C had the lowest plasticity (mean RDPI = 0.02; PI = 0.05); (2) stomatal length, stomatal density, SLA and vein density exhibited fivefold higher plasticity (RDPI = 0.10, 0.13, 0.12, 0.10, respectively; PI = 0.18, 0.26, 0.21, 0.22, respectively); (3) nitrogen content and trichome density had the highest (10-fold) plasticity (RDPI = 0.21, 0.36, respectively; PI = 0.32, 0.61, respectively). Polyploids and diploids, however, exhibited similar levels of plasticity for all seven traits (all *P* > 0.05 for RDPI and PI; Table S4).

### Is there a polyploid fitness advantage across diverse garden environments?

The main effect of ploidy level influenced plant fitness (*F* = 20.02, df = 1, *P* < 0.001; Fig. 3), after accounting for the significant negative effect of climatic niche distance (*F* = 71.54, df = 1, *P* < 0.001). Polyploids had significantly higher fitness than diploids at Corvallis (*t* = 3.86, *P* = 0.002) and Bend (*t* = 3.02, *P* = 0.011), and marginally higher at Newport (*t* = 1.97, *P* = 0.072), a pattern that refutes the ‘master-of-some’ strategy for polyploids or diploids. Fitness changed dramatically for both polyploids and diploids across the three gardens (garden effect: *F* = 563, df = 2, *P* < 0.001), contradicting fitness homoeostasis of the ‘jack-of-all-trades’ hypothesis but instead supporting the ‘jack-and-master’ hypothesis for polyploids.

**Figure 3:**
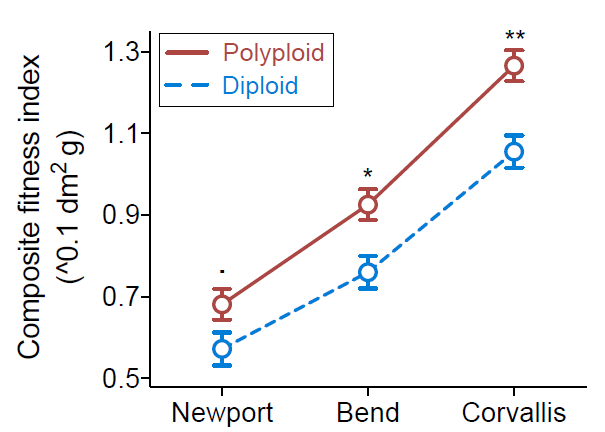
Fitness differences between diploid and polyploid *Fragaria*. The response variable was transformed (with a power parameter λ = 0.1) in the general linear mixed model. The *x*-axis shows garden locations. Error bars represent one standard error from the least-squares means. The dashed line depicts diploids, and solid line depicts polyploids. Significance levels: ^**^, *P* < 0.01; ^*^, *P* < 0.05; ?, *P* = 0.072.

### Is the polyploid fitness advantage conferred by trait means or trait plasticities?

For both diploids and polyploids, average fitness was influenced by trait means in four of the seven functional traits (i.e. stomatal length, SLA, vein density and trichome density; Fig. 4a, black symbols), with the strength often varying between ploidy levels. The trait mean of stomatal length had a significant positive effect on average fitness (Fig. 4a), indicating that plants with larger stomata had higher fitness, and the magnitude of this positive effect was similar between polyploids (β’ = 0.28, *P* < 0.001; Fig. 4b) and diploids (β’ = 0.28, *P* < 0.01). The mean of SLA also positively influenced average fitness (Fig. 4a), but the magnitude was stronger in polyploids (β’ = 0.27, *P* = 0.015) than diploids (β’ = 0.16, *P* = 0.074). While plants producing foliage of higher vein density and trichome density had lower fitness (Fig. 4a), these negative effects were especially strong in diploids (β’ = -0.25, *P* < 0.001; β’ = -0.30, *P* < 0.001, respectively) relative to polyploids (β’ = -0.08, *P* = 0.40; β’ = -0.10, *P* = 0.42, respectively).

**Figure 4:**
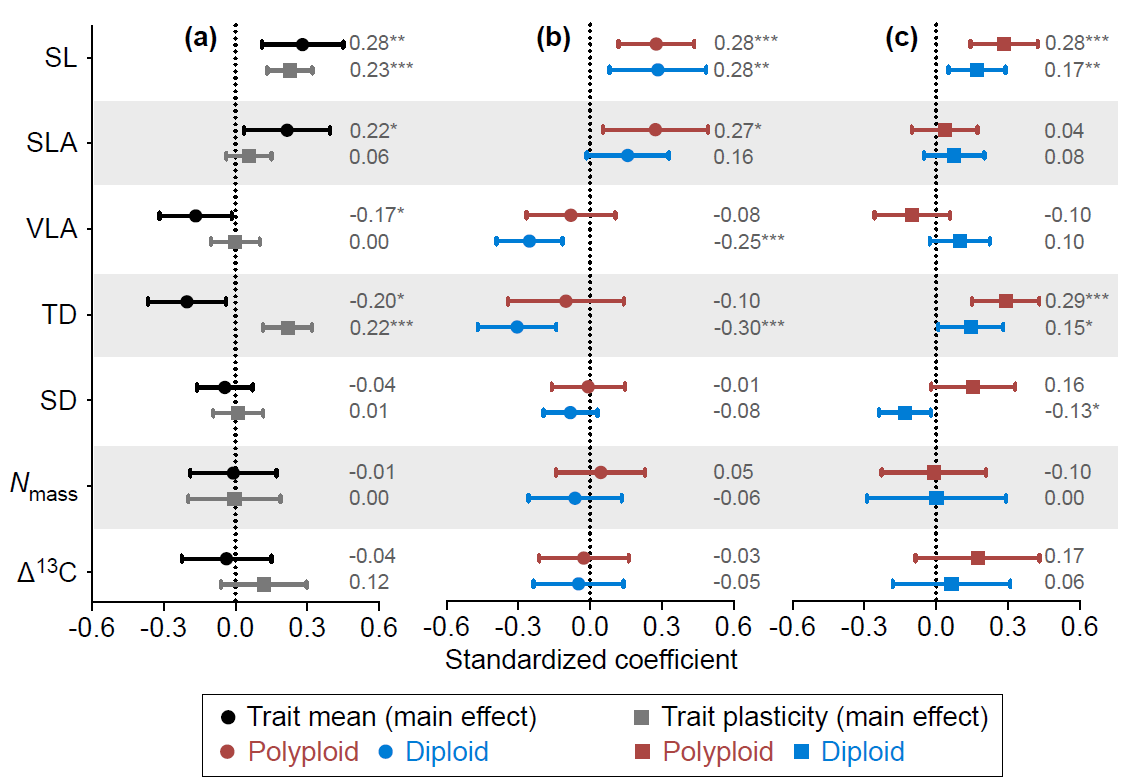
Trait means and trait plasticities predict average fitness of diploids and polyploids in the heterogeneous macroenvironment of this study. (a–c) Standardized regression coefficients of the main effects of trait means and plasticities (a), the ploidy-specific effects of trait means (b) and the ploidy-specific effects of trait plasticities (c), in models fitted separately for each functional trait. The average estimates are denoted by the symbols and the values to the right of error bars (95% confidence intervals). In (b) and (c), the polyploid estimate is above the diploid estimate for each trait. SL, stomatal length; SLA, specific leaf area; VLA, vein density; TD, trichome density; SD, stomatal density; *N*_mass_, nitrogen content; Δ^13^C, carbon isotope discrimination. Significance levels: ^***^, *P* < 0.001; ^**^, *P* < 0.01; ^*^, *P* < 0.05.

Trait plasticities had significant positive effects on average fitness for only two of the seven traits (Fig. 4a, gray symbols), and the strength of such adaptive plasticities in stomatal length and trichome density was nearly twofold higher in polyploids (β’ = 0.28 and 0.29, respectively; Fig. 4c) than diploids (0.17 and 0.15, respectively). Also noteworthy is that plasticity in stomatal density was maladaptive for diploids (β’ = -0.13, *P* = 0.019; Fig. 4c) but marginally adaptive for polyploids (β’ = 0.16, *P* = 0.084), despite the overall neutral effect on average fitness (Fig. 4a).

## DISCUSSION

Using a worldwide genotype collection of *Fragaria* grown in three different climatic regions (cool/coastal, temperate/valley, arid/montane), we derive general insight into the mechanisms underlying polyploid adaptation to heterogeneous environments. By comparing functional traits between ploidy levels, we find a cascading effect of genome size from the cellular level to trait level, which largely determines the phenotypic divergence between diploids and polyploids. In addition, the equivalency in trait plasticity between ploidy levels suggests that increased genomic redundancy does not necessarily translate into broader phenotypic amplitude in polyploids, as is often predicted (Stebbins 1971; Levin 1983). Nevertheless, our results show that both trait mean and trait plasticity contribute to higher polyploid fitness, and provide support for the ‘jack-and-master’ hypothesis for polyploid advantage over diploids in the genus *Fragaria*.

### Similar trait plasticity between diploids and polyploids

Our findings of equivalent phenotypic plasticity between diploids and polyploids contradict the long-held idea (Stebbins 1971; Levin 1983) that greater phenotypic plasticity in polyploids enables them to occupy broader ecological niches by expressing suitable phenotypes across a wider range of environments than diploids. Despite rich theories (Ramsey & Ramsey 2014), there have been few empirical evaluations of functional trait plasticity and polyploidy, and none as extensive as our study in terms of the geographic, genetic and phylogenetic diversity of the source material. A glasshouse experiment of trait plasticity in response to water variation (Manzaneda *et al.* 2015) revealed similar plasticity between annual allotetraploid *Brachypodium hybridum* and its diploid progenitor *B*. *distachyon* in stomatal conductance and Δ^13^C, although the diploid exhibited higher plasticity in photosynthetic rate. In response to nutrient variation (Sánchez Vilas & Pannell 2017), similar plasticity in SLA was found in glasshouse conditions between autotetraploid and allohexaploid cytotypes of the annual *Mercurialis annua*. For perennial allotetraploid and diploid *Centaurea stoebe* (Hahn *et al.* 2012), equivalent plasticity between ploidy levels was observed in all measured functional traits in response to water and nutrient variation in garden settings, and only a few traits exhibited higher plasticity in the polyploid in response to garden sites for one of two measuring occasions. These case studies, along with ours, draw the general picture of equivalency in plasticity of functional traits between diploids and polyploids. This pattern appears consistent across diverse plant genera, life-history strategies and environments, suggesting that it may well be the rule rather than the exception, at least for herbaceous polyploid plants.

There are several potential explanations for the lack of differentiation in trait plasticity between ploidy levels. First, genomic plasticity in gene expression and the resultant phenotypic variability (e.g. Gaeta *et al.* 2007) may quickly diminish during the course of polyploid formation, as a result of gene loss or silencing of duplicated copies (Adams & Wendel 2005), particularly for genes involved in essential biological processes such as photosynthesis (De Smet *et al.* 2013). Second, even given gene retention in polyploids, it is possible that only one copy responds to a specific selection agent of abiotic environments, such as in allotetraploid cotton (Liu & Adams 2007) where one copy of the alcohol dehydrogenase gene responds to cold stress and the other to water treatment owing to subfunctionalization of gene duplicates. Third, similar plasticity between ploidy levels may arise from biased gene expression towards one of the subgenomes in allopolyploids. Such subgenome dominance has been seen in both synthetic and wild polyploids (Jackson & Chen 2010; Grover *et al.* 2012), such as *Brassica rapa* (Cheng *et al.* 2016) and *Mimulus peregrinus* (Edger *et al.* 2017). Thus, linking gene expression of polyploids and diploids in common garden experiments to phenotypic plasticity will be critical for disentangling the mechanisms underlying similar trait plasticity between ploidy levels, as well as resolving when genomic plasticity (Adams & Wendel 2005; Leitch & Leitch 2008) is – or is not – correlated with phenotypic plasticity.

### Polyploid fitness advantage and its ecological mechanisms

Among the heterogeneous environments provided by our climatic gardens, polyploid *Fragaria* displayed the ‘jack-and-master’ strategy, showing higher fitness in each environment, and overall higher average fitness than diploids. Such polyploid fitness advantage has also been detected in the autotetraploids *Arrhenatherum elatius* (Petit & Thompson 1997) and *Dactylis glomerata* (Bretagnolle & Thompson 2001), and the allotetraploid *Centaurea stoebe* (Hahn *et al.* 2012). In a *Claytonia* complex (two 2*x*, one 4*x* and two 6*x* cytotypes) growing in California, one 6*x* cytotype possessed higher biomass than the others consistently across elevational gardens, albeit not for all polyploid cytotypes (McIntyre & Strauss 2017). In our study, polyploids performed as habitat generalists relative to diploids; yet, as they did not retain fitness homeostasis across climatic gardens, the ‘jack-of-all-trades’ hypothesis must be rejected. Plasticity in fitness (i.e. enhanced fitness under more favorable environment) is ubiquitous among the aforementioned polyploids showing fitness advantage over diploids, and has been seen to have similar magnitude between these polyploids and diploids (Petit & Thompson 1997; Bretagnolle & Thompson 2001; Hahn *et al.* 2012). Although diploid advantage was found in *Mercurialis annua* for aboveground biomass relative to the 6*x* cytotype in Spain (Buggs & Pannell 2007), it is unclear whether this advantage would hold when considering the belowground as well, as seen in this species between the 4*x* and 6*x* cytotypes in Morocco (Sánchez Vilas & Pannell 2017). While these previous studies and ours here often support the ‘jack-and-master’ hypothesis for polyploids, we nevertheless could not rule out the possibility that some diploid *Fragaria* may exhibit the ‘master-of-some’ strategy in environments beyond the climatic variation captured by this study, albeit our gardens are contained within the climatic niches of *Fragaria* species (Fig. S2) and niche distances were taken into account in our analyses. Thus, generalizing the adaptive strategies of polyploids and diploids will require not only genetically and geographically broad sampling of taxa as we have here, but also more diverse field environments than our study.

To our knowledge, this is the first study to explicitly link functional traits and plasticity to fitness differences between polyploids and diploids in the field. Although a study in 2*x* and 4*x Centaurea stoebe* (Hahn *et al.* 2012) also attempted to elucidate these mechanisms, their results focused on the contribution of trait plasticity (while controlling for trait mean), and found adaptive plasticity when considering both ploidies together as one group. Our study not only detected the importance of trait plasticity in determining polyploid and diploid fitness, but also revealed differential strength of adaptive plasticity (i.e. the slope of plasticity against fitness) between ploidy levels, which was higher in polyploids. Relative to trait plasticity, we found that functional trait divergence between polyploids and diploids, as a result of genomic changes in size and structure (Levin 1983; Balao *et al.* 2011), likely plays a more important role in determining fitness differentiation between ploidy levels, as more traits predict fitness in terms of trait means rather than plasticities. Also significant is that polyploids benefit from stronger positive fitness effects and weaker negative fitness effects of their functional traits, perhaps because their trait means are closer to optima than diploids in the experimental habitats.

In conclusion, the broad phylogenetic, genetic and geographic scope of this study provides the most robust evaluation to date of adaptive hypotheses for polyploid fitness advantage in changing environments, and elucidates functional trait divergence and adaptive plasticity as the underlying mechanisms. We emphasize that our findings are based on naturally occurring diploids and polyploids; as such, they reflect the ‘effective’ adaptions of polyploidy, resulting from the combined effects of polyploid formation and polyploidy-enabled establishment and divergence. Nevertheless, our results add significantly to the understudied ecological adaptations of polyploids, especially allopolyploids (Ramsey & Ramsey 2014), and offer important insight into the causes of evolutionary success of repeated and pervasive polyploidization (Van de Peer *et al.* 2017).

## AUTHORSHIP

TLA and AL designed research, RC managed garden establishment, and all authors contributed to design refinement. All authors collected data. NW performed data analyses. NW wrote the draft of the manuscript, and all authors contributed substantially to revisions.

## DATA ACCESSIBILITY STATEMENT

The data of genotype sampling, garden climate and plant functional traits and fitness are available from the Dryad Digital Repository.

## ACKNOWLEDGEMENTS

We are grateful to all the collectors of wild strawberry genotypes. We thank K. Schuller, T. Jennings, C. Poklemba, P. Krabacher, M. Surplus, A. Kessler and L. Longway for assistance with genotype and clone cultivation, common garden establishment and management, and fitness measurements, A. Freundlich, E. Jiang, C. Rumrill, R. Babu, I. Alagar, S. Barratt-Boyes and Z. Du for assistance with leaf collection and functional trait measurements, and C. Muir for help with impression technique of leaf stomata. We acknowledge the University of Pittsburgh glasshouses, Oregon State University glasshouses and Hatfield Marine Science Center, and USDA Forest Service Region 6 Bend Seed Extractory for logistic support, and the members of the Ashman, Liston, and Cronn labs for discussion. This work was supported by the National Science Foundation (DEB 1241006 to TLA; DEB 1241217 to AL).

